# Feasibility and promise of circulating tumor DNA analysis in dogs with naturally-occurring sarcoma

**DOI:** 10.1101/2020.08.20.260349

**Authors:** Patricia Filippsen Favaro, Samuel D. Stewart, Bradon R. McDonald, Jacob Cawley, Tania Contente-Cuomo, Shukmei Wong, William P.D. Hendricks, Jeffrey M. Trent, Chand Khanna, Muhammed Murtaza

## Abstract

Comparative studies of naturally-occurring canine cancers have provided new insight into many areas of cancer research. The inclusion of pet dogs in the development and validation of circulating tumor DNA (ctDNA) diagnostics may be uniquely informative for human translation for many reasons, including: high incidence of certain spontaneous cancers, repeated access to blood and tumor from the same individuals during the course of disease progression, and molecular heterogeneity of naturally-occurring cancers in dogs. Here, we present a feasibility study of ctDNA analysis performed in 9 healthy dogs and 39 dogs with either benign or malignant splenic tumors (hemangiosarcoma) using shallow whole genome sequencing (sWGS) of cell-free DNA. To enable detection and quantification of ctDNA using sWGS, we adapted two informatic approaches and compared their performance for the canine genome. At presentation, mean ctDNA tumor fraction in dogs with malignant splenic tumors was 11.2%, significantly higher than dogs with benign lesions (3.2%; p 0.001), achieving an AUC of 0.84. ctDNA tumor fraction was 14.3% and 9.0% in dogs with metastatic and localized disease, respectively although this difference was not statistically significant (p 0.227). In paired analysis, ctDNA fraction decreased from 11.0% to 7.9% after resection of malignant tumors (p 0.047). Our results demonstrate that ctDNA analysis is feasible in dogs with hemangiosarcoma using a cost-effective approach such as sWGS. Future studies are underway to validate these findings, and further evaluate the role of ctDNA to assess burden of disease and treatment response during drug development.

## Introduction

Comparative studies of naturally-occurring canine cancers provide a unique opportunity to advance the care of human cancer patients by answering questions about cancer biology, genomics, diagnosis, and therapy, that cannot be answered in conventional preclinical cancer models or in human clinical trials (Paoloni and Khanna 2008; Paoloni et al. 2014; Gustafson et al. 2018). The analysis of somatic mutations in canine cancers can yield novel insights into pathogenesis of the disease (Lorch et al. 2019). Exome sequencing of canine splenic hemangiosarcoma (HSA) identified somatic mutations in well-known driver genes of human cancer (Wang et al. 2017). Unlike most preclinical models, the molecular heterogeneity seen in spontaneous cancers in dogs mimics human cancer heterogeneity and can serve as a unique model system for development of novel therapeutic and diagnostic approaches.

Analysis of circulating tumor DNA (ctDNA) in dogs with cancer may help further our understanding of ctDNA biology due to: 1) the high incidence of spontaneous cancer in dogs, including those that are rare in humans (such as sarcomas); 2) the opportunity to collect serial blood samples of sufficient volume compared to conventional preclinical models and to collect concurrent samples of tumor and normal tissue, using clinical annotation methods commonly employed in human oncology (such as tumor grade, and patient stage); 3) comparable relative sizes of tumors between dogs and humans; and 4) the ability to optimize and compare sample collection and data analysis approaches in the same individual. However, published experience with ctDNA analysis in dogs is currently sparse. Nonetheless, limited and small-scale published studies have suggested total cell-free DNA concentrations are higher in dogs with malignant tumors compared to those with benign disease or healthy controls (Schaefer et al. 2007; Beffagna et al. 2017; Tagawa et al. 2020). In a recent study of canine mammary carcinoma, ctDNA detection was demonstrated using digital PCR assays specific to somatic genomic rearrangements in 4 dogs (Beck et al. 2013). In another study, ctDNA detection was demonstrated in 8 of 11 dogs with urothelial carcinoma using a real-time PCR assay for a recurrent somatic mutation in BRAF (Tagawa et al. 2020). Earlier, we demonstrated detection of ctDNA in 2 of 6 dogs with pulmonary adenocarcinoma using a digital PCR assay for a recurrent somatic mutation in ERBB2 (Lorch et al. 2019). Hence, published demonstrations of ctDNA analysis are limited to a few dogs and have largely relied on mutation and locus-specific assays. Mutation-specific assays have limited widespread applicability for ctDNA analysis in dogs or humans because they either rely on highly recurrent single mutations or require prior analysis of the tumor sample to identify patient-specific mutations. In this study, we have evaluated an alternative approach through the use of shallow Whole Genome Sequencing (sWGS) for ctDNA analysis of canine cancer, which relies on direct genome-wide assessment in plasma DNA for copy number aberrations.

Canine HSA is a relatively common canine cancer with strong clinical similarities to human angiosarcoma. Both cancers harbor structurally complex genomes that are not associated with recurrent point mutations, limiting the utility of single mutation ctDNA assays. Hemangiosarcoma commonly develops in the spleen of dogs and metastasizes to distant organs prior to or early after initial diagnosis. These splenic lesions are not often discovered until they rupture and the dog presents to the veterinary hospital with acute abdominal hemorrhage, requiring emergent surgery. This clinical presentation is very similar to dogs with benign tumors of the spleen or other causes of splenic rupture. Histopathological confirmation of the diagnosis for these lesions cannot usually be achieved until several days after emergency surgeries are performed, highlighting the challenges that pet owners face when choosing to pursue aggressive emergent surgical care and treatment for their dog with an unclear long-term prognosis. An opportunity exists to study and translate methods for ctDNA analysis in this naturally-occurring cancer model, to ask if ctDNA levels can distinguish benign from malignant tumors, to understand if ctDNA levels are related to burden of disease at presentation (such as localized vs. metastatic cancer), and if serial changes in ctDNA levels are effective surrogates for assessment of treatment response or disease progression. These investigations will be relevant to the application of ctDNA analysis within human oncology and allow for optimization of diagnostic methods and technologies. Here, we present a feasibility study evaluating the potential for ctDNA analysis in dogs with cancer using sWGS to detect ctDNA in dogs presenting with benign or malignant splenic tumors. We hope our findings and adapted informatics tools will help expand the inclusion of dogs with naturally-occurring cancers as models for optimal ctDNA assay development and translation, and aid drug development efforts.

## Results

### Cell-free DNA analysis in healthy dogs

In order to establish the feasibility of cell free DNA (cfDNA) analysis in canines, we obtained plasma samples from 9 healthy dogs including five females and four males. The mean cfDNA concentration was 0.39 ng/ml of plasma (sd 0.38 ng/ml). We prepared sequencing libraries with 1-30 ng of DNA input and generated a mean of 20.0 million sequencing read pairs per sample. We achieved a mean unique depth of 0.45x per sample (sd 0.15). Following alignment to the dog genome CanFam3.1, we determined the modal fragment size of 165.6 base pairs (bp), comparable to the known modal fragment size of 166 bp in human cfDNA (Jiang et al. 2015; Murtaza and Caldas 2016)(Figure 1, Table S1).

**Figure 1.**
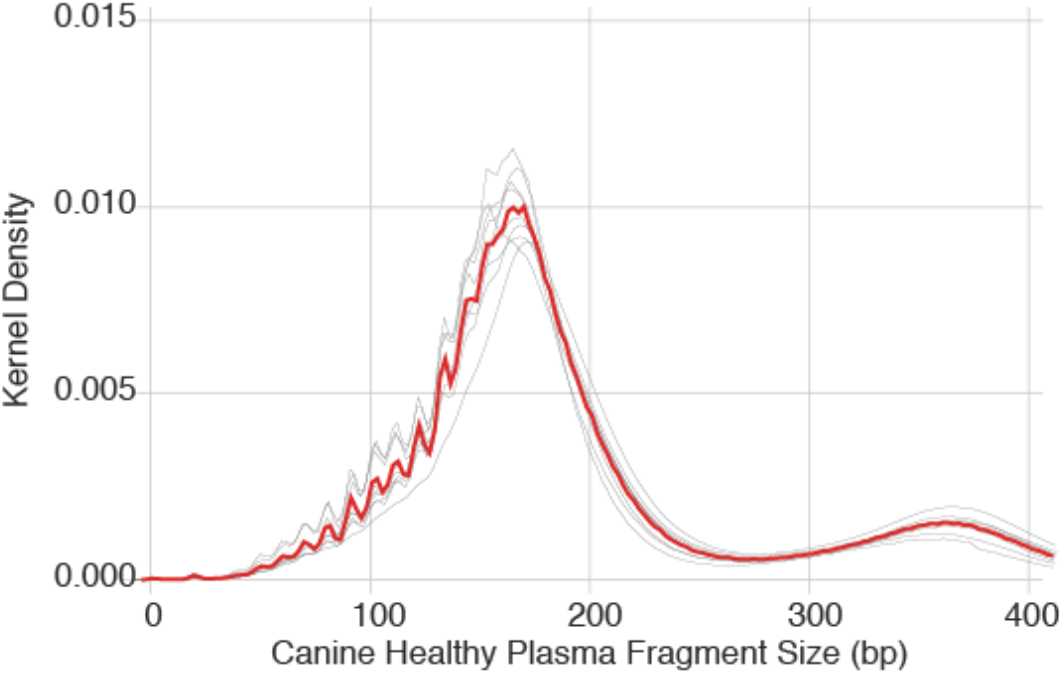
Plasma cell-free DNA fragment size (base pairs) in nine healthy dogs. Individual samples are show in grey, all samples combined are shown in red.

### Clinical cohort of dogs with splenic lesions

To determine the feasibility of cfDNA analysis in dogs, we conducted a prospective clinical trial of dogs presenting with hemo-abdomen secondary to presumed splenic rupture. The description of this clinical cohort of dogs, including diagnoses and perioperative outcomes, has been recently reported (Stewart et al. 2020). Samples from 39 dogs from this cohort were evaluable for cfDNA analysis. Twenty five dogs (64.1%) were classified as having malignant splenic neoplasms and 14 were diagnosed with benign splenic lesions (35.9%). Among the 25 dogs that were diagnosed with malignant tumors, 23 (92%) were diagnosed with HSA and 2 were diagnosed with stromal sarcoma. Benign complex nodular hyperplasia represented 78.6% (11/14) of the benign lesions, while complex hyperplasia with hematoma, hematoma alone and myelolipoma represented 7.14% each (1/14 each). Two dogs (5.13%) had concomitant benign lesions and malignant tumor nodules in different portions of the spleen. These were grouped and analyzed as dogs with malignant disease in the current analysis.

### Comparison of informatic approaches for sWGS

In order to investigate the feasibility of cfDNA analysis in dogs using sWGS, we adapted two published informatic approaches, ichorCNA (Adalsteinsson et al. 2017) and PlasmaSeq (Farris and Trimarchi 2013), to enable their application to non-human genomes. Both tools infer the presence of tumor-derived copy number alterations (CNAs) using read depth in large, non-overlapping genomic bins (windows). Bins are then grouped into segments with the same copy number status. ichorCNA uses fixed-size bins while PlasmaSeq uses a fixed total number of bins and the boundaries are adjusted so that each bin contains approximately the same number of mappable bases. ichorCNA uses the magnitude represented by the detected CNAs to directly infer the fraction of tumor DNA found in blood. The canine patient samples were analyzed using 500 kb bins for ichorCNA. PlasmaSeq analysis was run using 5,500 total bins, with median size of 392 kb (sd 42 kb). Across all plasma samples from dogs with splenic lesions, the total number of segments was significantly higher for PlasmaSeq, with an average of 72.6 segments per sample versus 50.0 segments per sample for ichorCNA. PlasmaSeq had a larger proportion of segments with less than 25 bins and more than 150 bins, while ichorCNA had a higher proportion of segments with 50-100 bins (Figure 2a). We also quantified the amount of variation in read depth (log2 values) within bins assigned to a given copy number segment. The median standard deviation within each segment for ichorCNA and PlasmaSeq were 0.0606 and 0.0569 respectively, (Figure 2b, p 0.004 Mann-Whitney U). Although distinct, we found the two adapted approaches to be equally useful for cfDNA analysis in dogs.

**Figure 2.**
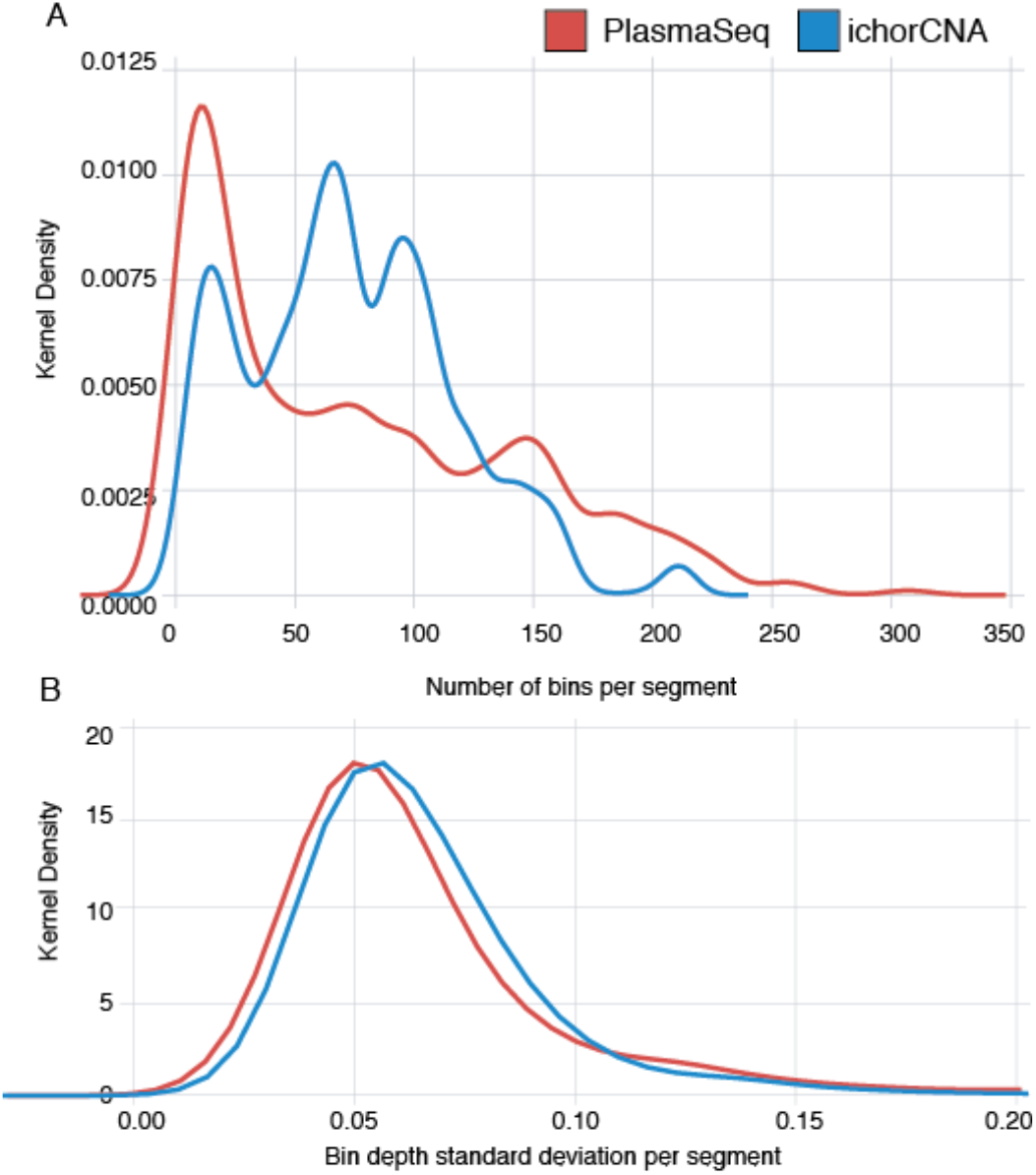
PlasmaSeq infers a larger proportion of segments with less than 25 bins and more than 150 bins, although the depth variation within segments is approximately the same between PlasmaSeq and ichorCNA. A: The distribution bin counts in each segment across all samples using each tool. B: The distribution of log2 standard deviations across all bins assigned to the same segment.

### Analysis of total cell-free and circulating tumor-specific DNA

At presentation prior to any intervention, mean concentration of total cfDNA in plasma samples was 10.6 ng/ml and 20.3 ng/ml in dogs with benign and malignant lesions, respectively (p 0.400). Total cfDNA concentration after surgery was significantly lower than baseline in both groups (benign p 0.008 and malignant p 0.017). Within dogs with malignant lesions, mean total cfDNA concentration was 20.9 ng/ml and 19.5 ng/ml in dogs with localized and metastatic disease, respectively (p 0.830).

ctDNA tumor fractions reported here were measured using an adapted version of ichorCNA. Mean tumor fractions prior to surgery were 3.2% (sd 3.4) and 11.2% (sd 9.1) in dogs with benign and malignant tumors, respectively (p 0.001; Figure 3, Table S2). Within dogs with malignant tumors, mean ctDNA fractions in dogs with localized and metastatic disease were 9.0% and 14.3% respectively, although this difference was not statistically significant (p 0.227). Following splenectomy, patients with malignant tumors showed a significant decrease in ctDNA levels (paired p 0.047). ctDNA was detected above the previously reported limit of detection for ichorCNA of 3% tumor fraction in 20/21 dogs with malignant tumors with a pre-treatment sample available (95.2%). Using tumor fractions in ctDNA, we were able to distinguish blood samples from dogs with benign and malignant tumors with an area under the ROC curve of 0.84 (Figure 4). Interestingly, when corresponding tumor biopsies were analyzed using sWGS, we observed mean tumor fractions of 4.4% (sd 4.9) and 10.1 % (sd 12.9) for benign and malignant tumors, respectively, highlighting the challenges of obtaining high-cellularity splenic tumor biopsies and establishing tissue diagnoses in dogs, and the additional value potentially offered by ctDNA analysis.

**Figure 3.**
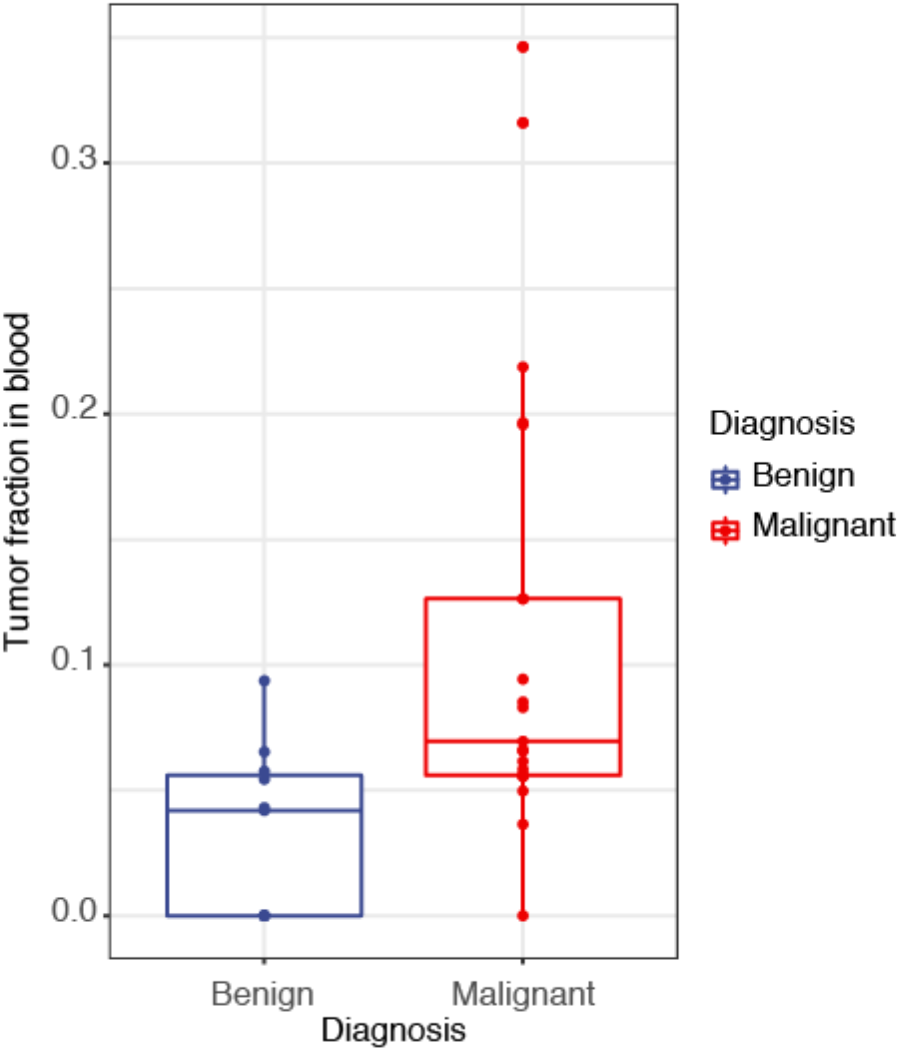
cfDNA tumor fraction (TF) for benign vs malignant tumors in the baseline sample cohort (p = 0.001).

**Figure 4.**
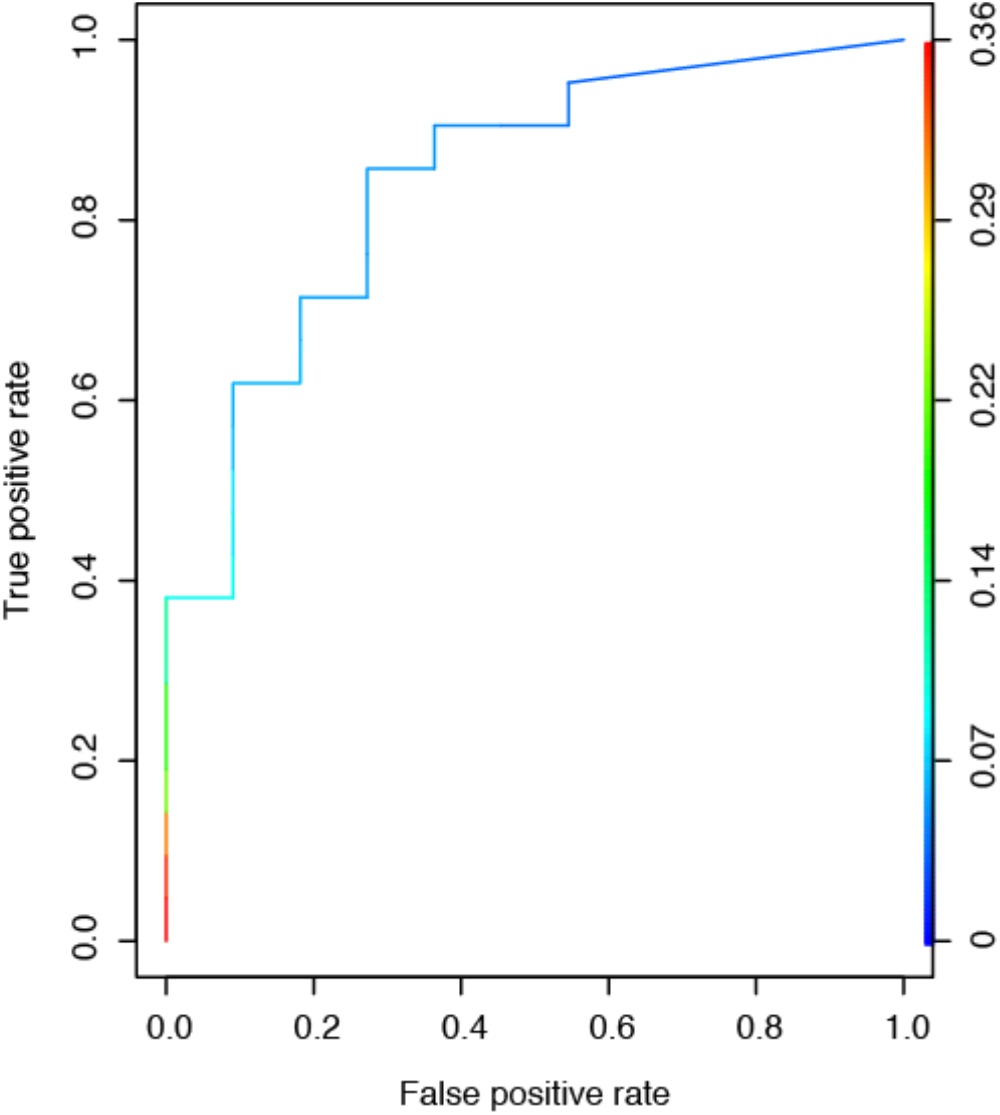
Receiver operating characteristic (ROC) curve for tumor fraction detection in cell free DNA from plasma from 32 dogs with benign and malignant tumors before surgery, with AUC = 0.84.

We found a statistically significant bias in the distribution of somatic copy number changes across chromosomes, although this bias was not consistent across tools. Chromosomes 8, 11, 14, and 26 were more likely to contain copy number gains called by PlasmaSeq, while chromosomes 5 and 34 were more likely to have gains called using ichorCNA (Table 1).

**Table 1.**
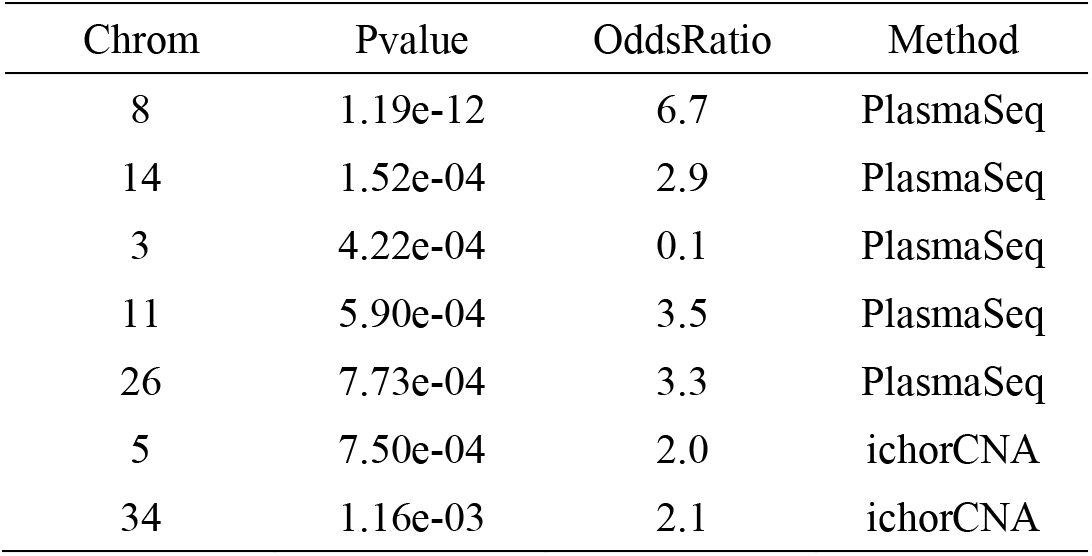
Copy number variation odds ratio of the chromosomes in the 39 canine patients with splenic lesions. P values and odds ratios were calculated using Fisher’s Exact Test.

| Chrom | Pvalue | OddsRatio | Method |
| --- | --- | --- | --- |
| 8 | 1.19e-12 | 6.7 | PlasmaSeq |
| 14 | 1.52e-04 | 2.9 | PlasmaSeq |
| 3 | 4.22e-04 | 0.1 | PlasmaSeq |
| 11 | 5.90e-04 | 3.5 | PlasmaSeq |
| 26 | 7.73e-04 | 3.3 | PlasmaSeq |
| 5 | 7.50e-04 | 2.0 | ichorCNA |
| 34 | 1.16e-03 | 2.1 | ichorCNA |

In cases where corresponding tumor DNA and cfDNA samples both showed adequate tumor fraction, copy number aberrations observed were largely concordant. For example, in one dog with metastatic malignant hemangiosarcoma, copy number aberrations observed in tumor biopsy are also observed in blood samples obtained on days 0, 16 and 19 after splenectomy, using both informatic approaches (Figure 5).

**Figure 5.**
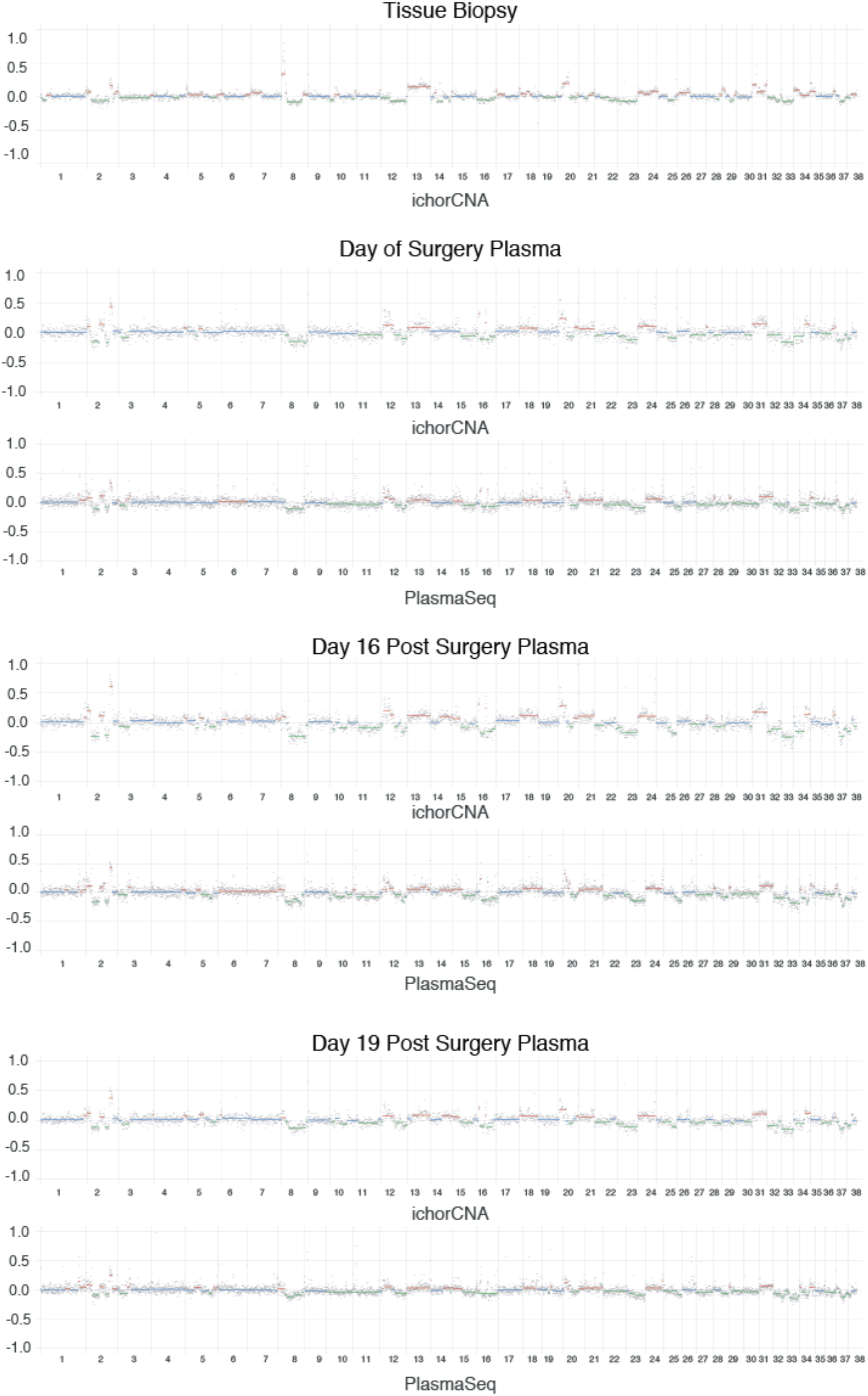
Genome-wide copy number variation plots from the metastatic HSA canine patient, dog no. 26, representing the plasma and tumor biopsy sWGS from day of surgery, and plasma sWGS from days 16 and 19, both after the splenectomy procedure.

Follow-up blood samples from two dogs classified as having a) benign complex nodular hyperplasia and b) complex nodular hyperplasia with hematoma, had unexpectedly high tumor fractions in plasma (21.4% and 46.9%). However, these observations were made at 392 and 345 days after surgery, respectively. In both dogs, earlier samples had ctDNA levels consistent with those observed in dogs with benign disease (5.4% and 4.8%). The source of these elevated ctDNA levels observed later in follow-up is unclear. In these older clinical patients, it is possible that elevated ctDNA levels almost a year after initial diagnosis are a result of clinically silent unrelated cancers. Indeed, post hoc analysis revealed new neurologic clinical signs that were localized to the right cerebral cortex 415 days following surgery in one of the two dogs suggestive of distinct intracranial neoplasia. However, further investigation was not pursued to confirm this diagnosis. There were no clinical signs suggestive of neoplasia in the second dog. Nonetheless, the presence of indolent neoplasia (such as indolent lymphoma) in this clinical cohort of older large breed dogs cannot be ruled out.

## Discussion

ctDNA analysis in pet dogs with cancers provides an opportunity to develop and validate methods, and to optimize pre-analytical conditions for molecular and computational analysis of liquid biopsies. Analysis of naturally-occurring cancers with similar complexities of cancer heterogeneity as human patients can potentially generate novel insights and allow rapid translation back to human oncology. However, published experience with ctDNA analysis in canine oncology has been very limited. In the current study, we demonstrate that sWGS is useful as a tumor-independent approach for ctDNA detection in dogs with cancer. We validate the application of informatic methods adapted for the canine genome and make these adapted versions available for future use to accelerate the development and deployment of ctDNA analysis in dogs (Farris and Trimarchi 2013; Adalsteinsson et al. 2017). We provide proof-of-principle results suggesting potential utility of ctDNA analysis using sWGS in dogs presenting with structurally complex cancers to specifically distinguish malignant cancers from benign disease and as a potential surrogate of disease burden. The analysis of ctDNA levels in the same dog before and after resection of the splenic tumors yielded significant differences in ctDNA levels, whereas comparison of dogs with localized versus metastatic disease (noted at the time of surgery) were not statistically distinct. Given the relatively limited size of our cohort, comparison of ctDNA levels with metastatic disease burden requires further analysis in a larger cohort of dogs. In future studies, we plan to include measurement of tumor volumes to account for quantitative differences in burden of metastatic disease and size of primary tumors, and how these may correlate with observed differences in ctDNA levels. Furthermore, we plan to perform paired comparisons of sample processing workflows and analytical approaches to optimize measurement of ctDNA levels as a surrogate for disease burden. With further clinical validation in larger studies, ctDNA analysis can aid diagnostic dilemmas such as those investigated here and may be included in studies that can explore the feasibility and value of early diagnostic approaches, better prognostication, and improved treatment monitoring for rapid translation to human oncology.

This feasibility study has several limitations. ctDNA studies in human oncology have largely come to rely on gold standard pre-analytical processing focused on rapid separation and adequate processing of plasma samples. In on-going studies, we have ensured implementation of standard processing protocols to isolate plasma rapidly after venipuncture. In future work, ctDNA analysis in dogs may provide an opportunity to evaluate and optimize pre-analytical factors using paired tumor samples and multiple blood samples from cancer patients, such as blood collection tubes, DNA extraction methods, and storage conditions. An additional limitation was our observation of significantly elevated levels of ctDNA detectable in two dogs with benign cancers in follow-up blood samples. These dogs were shown to have benign splenic lesions at the time of initial diagnosis and surgery but hematological or other malignancies present concomitantly or arising later may only become apparent during sufficient clinical follow-up. In one of these dogs, concomitant neoplasia of the brain was suspected, but not confirmed, based on the subsequent development of clinical signs suggestive of a space occupying mass. Given the clinical characteristics of this cohort of larger older dogs, we speculate that a non-splenic cancer, such as indolent lymphoma, may explain these findings.

In summary, our results demonstrate that ctDNA analysis is a promising approach for improving diagnostics in veterinary oncology. In addition, ctDNA analysis of naturally-occurring cancers in dogs can enable further optimization for diagnostic applications in human oncology including noninvasive tumor genotyping, early detection, and monitoring of treatment response.

## Methods

### Ethics statement

Blood and tissue samples were collected with the approval and consent of the canine patient owners. The study was launched following approval from the animal care and use committee convened by Animal Clinical Investigation (Chevy Chase, MD).

### Sample collection

Blood samples were collected from the jugular or cephalic vein from 39 canine patients that were presenting acute hemoperitoneum secondary to splenic neoplasm rupture. All dogs underwent preoperative staging, the results of which have been previously reported (Stewart et al. 2020). The whole blood was transferred to Cell Free DNA BCT Streck tubes (Streck, Inc.), processed as per manufacturer’s instructions and the collected plasma and buffy coat were stored at −80° C. The blood sample collections were performed prior to the splenectomy procedure and during the subsequent clinical visits. Biopsy sections of 1cm x 1cm x 1cm were collected from three distinct regions of the spleen: primary tumor site, tumor periphery and visually normal splenic tissue, and were stored in 10% formalin at 4° C for histopathological analysis. Additional biopsy sections were collected from the same regions and at two distinct areas of the tumor were placed in Allprotect Tissue Reagent (Qiagen, Hilden, Germany) and stored at 4° C for DNA extraction and evaluated histopathologically. Through the clinical and histopathological examinations, the patients were diagnosed with malignant splenic neoplasms (n = 25) and benign splenic neoplasm (n = 14). All histopathologic diagnoses were verified through post hoc medical record reviews and the clinical data was locked to confirm the diagnoses. Blood samples were also collected in Cell Free DNA BCT Streck tubes (Streck, Inc.) from nine healthy dogs and the plasma frozen at −80° C. The plasma samples were used for cfDNA extraction and the tissue biopsy for total DNA extraction, which were used for shallow whole genome sequencing.

### DNA extraction from tumor, white blood cells and plasma

Cell-free DNA extractions from 1-4 ml of plasma were performed with MagMAX Cell-Free DNA Isolation Kit (Thermo Fisher Scientific, Austin, TX) or the QIAmp Circulating Nucleic Acid Kit (Qiagen, Hilden, Germany), as per manufacturer’s instructions. The tissue biopsy (30 mg) was rinsed in 1X phosphate buffered saline, homogenized with Bullet Blender Bead Lysis Kit (NextAdvance, Troy, NY) and the supernatant passed through QIAshredder columns (Qiagen, Hilden, Germany) before been used for tumor DNA extraction using the Allprep DNA/RNA/miRNA Universal Kit (Qiagen, Hilden, Germany), according to the manufacturer’s instructions. All extracted DNA were stored at −20° C until use. The cfDNA and tumor DNA concentration, integrity and purity were assessed with the Bioanalyzer (Agilent Technologies, Santa Clara, CA), Qubit 2.0 Fluorometer (Thermo Fisher Scientific, Austin, TX) and 4200 TapeStation genomic DNA assay (Agilent Technologies, Santa Clara, CA). DNA from the white blood cells were extracted from 200 μl of buffy coat with the DNeasy Blood Tissue Kit (Qiagen, Hilden, Germany), according to the manufacturer’s instructions, and stored at −20° C until further processing.

### cfDNA and tumor DNA next generation sequencing

The cfDNA library construction was performed with the SMARTer ThruPLEX Plasma-Seq Kit and DNA HT Dual Index Kit (Takara Bio USA, Mountain View, CA), as per manufacturer’s instructions. Sequencing libraries were purified with SPRI magnetic beads (Beckman Coulter, Brea, CA). Library sizes and concentrations were measured using a genomic DNA assay on the TapeStation 4200 (Agilent Technologies, Santa Clara, CA). Tumor DNA was fragmented to a target size of 200 bp on E220 Focused-ultrasonicator (Covaris, Woburn, MA). 20 ng of sheared DNA was used for the library construction with ThruPLEX DNA Seq Kit and DNA HT Dual Index Kit (Takara Bio USA, Mountain View, CA). Plasma and tumor DNA libraries were sequenced using paired-end 50 bp reads generated on the NovaSeq 6000 Sequencing System (Illumina, San Diego, CA).

### Somatic copy number analysis

Raw sequence data was converted to fastqs using illumina’s bcl2fastq v2.20.0.422. Reads were trimmed using fastp v0.2 (Chen et al. 2018) and then aligned to the canFam 3.1 (Hoeppner et al. 2014) genome assembly using bwa-mem. WGS bams from nine healthy canine plasma cfDNA samples were used as controls for ichorCNA and PlasmaSeq analysis of large-scale copy number changes. GenMap (Pockrandt et al. 2019) was used to calculate mappability on the canFam 3.1 genome assembly. PlasmaSeq analysis was conducted using our implementation of the algorithm using the julia programming language v1.1. Usable bases were identified using the mappability data, and boundaries for the fixed 5,500 bins were selected to reduce the variability of usable bases across all bins. The number of bins was selected to be approximately the same as the number of bins used by ichorCNA. For relative bin depth normalization, bams from the nine healthy control samples were merged. Read depths for PlasmaSeq bins were calculated using bedtools v2.28.0 (Quinlan and Hall 2010).

ichorCNA analysis was conducted using a modified version of ichorCNA v0.3.2. Functions that standardized chromosome names for human genomes were removed, as they caused errors with unexpected chromosome names. A canFam 3.1 panel of normals was generated using the nine healthy control samples with 500 kb bins. Mappability and GC content calculations were performed as for PlasmaSeq. HMMcopy (Lai et al. 2020) was used to calculate read depths per bin. Initial normal proportion range was set to [0.7,0.8,0.9,0.95,0.99]. Ploidy was fixed at 2, with a max copy number of 3 and subclone state options of [1,3]. Without these restricted parameter ranges, high tumor fraction inferences with no obvious copy number changes and inferred ploidy of 3 were very common across samples.

### Statistical analysis

Patient groups were compared using non-parametric tests including Kruskal-Wallis and Wilcoxon Rank-Sum tests using the R package *stats*, and plots were prepared using the packages *ggpubr, magrittr, ggplot2, ggsci* and *scales*. Specificity, sensitivity and test performance were calculated with the packages *ROCR* and *cvAUC*. All means, standard deviations and confidence intervals were calculated with the R package *stats*. The Fisher Exact Test was performed with the julia package *Hypothesis Tests*.

## Data Access

Upon manuscript acceptance, the sequencing data generated in this study will be submitted to the Sequence Read Archive (SRA) and the accession number will be updated. Adapted versions of ichorCNA and PlasmaSeq will be deposited to github and accession numbers will be updated here.

## Acknowledgments

We would like to thank pet families who participated in this study.

## Author Contributions

CK, MM, JT and WH conceptualized and designed the study. PFF, BRM, TCC and MM developed methods. SDS, JC and CK designed and conducted prospective clinical studies. PFF, TCC, and SW generated data. BRM and PFF analyzed sequencing data. PFF, SDS, BRM, CK and MM interpreted data. PFF, SDS, BRM, CK and MM wrote the paper with assistance from JC, TCC, SW, WH and JT. All authors approved the final manuscript.

